# Active exploration of a novel environment drives the activation of the hippocampus-amygdala complex of domestic chicks

**DOI:** 10.1101/2022.02.24.481770

**Authors:** Anastasia Morandi-Raikova, Uwe Mayer

**Affiliations:** Center for Mind/Brain Sciences (CIMeC), University of Trento, Piazza Manifattura 1, I-38068, Rovereto (TN), Italy

**Author notes:** Corresponding author: Asst. Prof. Uwe Mayer, Center for Mind/Brain Sciences, University of Trento, Piazza Manifattura 1, I-38068, Rovereto (TN).

**Keywords:** avian hippocampal formation, place cells, immediate early genes, spatial map, c-Fos, nucleus taeniae of the amygdala

## Abstract

In birds, like in mammals, the hippocampus critically mediates spatial navigation through the formation of a spatial map. This study investigates the impact of active exploration of a novel environment on the hippocampus of young domestic chicks, during formation of a new spatial map. Chicks that were free to actively explore the novel environment exhibited a significantly higher neural activation (measured by c-Fos expression), compared to those that passively observed the novel environment from a restricted area. The difference was limited to the anterior and the dorsolateral parts of the intermediate hippocampus. Furthermore, the nucleus taeniae of the amygdala showed a higher c-Fos expression in the active exploration group than the passive observation group. In both brain regions, brain activation correlated with the number of locations that chicks visited during the test. This suggest that the increase of c-Fos expression in the hippocampus is related to increased firing rates of spatially coding neurons. Furthermore, our study indicates a functional linkage of the hippocampus and nucleus taeniae of the amygdala in processing spatial information. Overall, with the present study, we confirm that, in birds like in mammals, exploration of novel environments activates hippocampus, which is likely related to the formation of new spatial representations.

**Summary statement:** Active exploration of a novel environment induces stronger activation of hippocampus and taeniae of domestic chicks than pure visual, passive exploration from a restricted area.

## INTRODUCTION

The neuroanatomical organization of the hippocampus (Hp) homologues differs dramatically between birds and mammals (Striedter, 2016). Despite these differences, in both these taxa the Hp supports similar cognitive functions. One example is represented by spatial navigation, which depends on the hippocampus in mammals (O’Keefe and Dostrovsky, 1971; Morris et al., 1982; Lavenex and Lavenex, 2009), birds (Fremouw et al., 1997; Bingman et al., 1984; Mayer et al., 2013; Bingman and Muzio, 2017; Morandi-Raikova and Mayer, 2021), reptiles (Day et al., 2001; López et al., 2003), amphibians (Sotelo et al., 2016) and fish (Rodrìguez et al., 2002). However, the extent to which these functions share similar neural mechanisms across vertebrate species needs further investigations.

Animal navigation is mainly based on ‘internal’ cognitive maps (Tolman, 1948). At least in mammals, the cognitive maps are formed through the critical contribution of hippocampal place cells (O’Keefe and Dostrovsky, 1971; O’Keefe and Conway, 1978). These cells exhibit spatially localized increase in firing rates when an animal occupies a specific field in an environment. Since the seminal discovery of place cells, many other spatially responsive cells have been also described in mammals (Moser et al., 2017; Poulter et al., 2018). Head direction cells (Taube et al., 1990a, b), grid cells (Sargolini et al., 2006) border cells (Sargolini et al., 2006), speed cells (Kropff et al., 2015) and vector trace cells (Poulter et al., 2021) all contribute to the formation of a cognitive map in mammals. In birds too, hippocampal place cells (Payne et al., 2021) and head direction cells (Ben-Yishay et al., 2021; Takahashi et al., 2022) have been recently found. Payne et al., (2021) found place cells in two different species of birds: tufted titmice (*Baeolophus bicolor*) and zebra finches (*Taeniopygia guttata*). Interestingly, the spatially responsive cells observed in these two bird species were located predominantly in the anterior hippocampus. The density of place cells decreased along the anterior-posterior axis. In rodents, place cells follow a similar gradient along the dorsoventral axis (Jung et al., 1994). The avian subregions along the anterior-posterior axis might thus be equivalent to the hippocampal regions along the dorsoventral axis in mammals (Payne et al., 2021). Indeed, following the same assumption, Agarwal et al., (2021 preprint) recorded place cells within the anterior hippocampus in freely flying barn owls (for earlier studies with pigeons see also Bingman et al., 2003; Hough and Bingman, 2004; Siegel et al., 2005, 2006; Kahn et al., 2008). Overall, these studies suggest that spatial processing in birds and mammals is based on similar mechanisms. Moreover, a higher number of spatially coding cells can be expected in the anterior region of bird’s hippocampal formation.

The importance of the avian hippocampus for spatial navigation has been often addressed using immediate early genes (IEGs) as neural activity markers (Smulders and DeVoogd, 2000; Bischof et al., 2006; Mayer et al., 2010; Mayer and Bischof, 2012; Grella et al., 2016; Sherry et al., 2017). As an alternative to electrophysiology, the use of IEG products offers a practical approach to investigate the activation of entire neural ensembles (Lanahan and Worley, 1998). Neural IEG expression rapidly increases in response to trans-synaptic signalling between neurons and the resulting genomic response is closely associated with neuronal plasticity (Jones et al., 2001; Guzowski, 2002; Barry and Commins, 2011). Using the IEG product c-Fos to study hippocampal activity in domestic chicks (*Gallus gallus domesticus*), it has been shown that this structure has many similar functions to its mammalian counterpart. Like in mammals, chicks’ hippocampus is sensitive to environmental boundaries. It shows high levels of c-Fos during navigation by the geometrical shape of the environment (Mayer et al., 2016) and responds to changes in the shape of the environment (Mayer et al., 2018). Furthermore, chicks’ hippocampus process spatial relational information, showing high expression of c-Fos during navigation in relation to free-standing objects (Morandi-Raikova and Mayer, 2021).

More importantly for the current study, in chicks, like in mammals, exposure to novel environments induces high levels of c-Fos expression within the hippocampus (Morandi-Raikova and Mayer, 2020; for mammals see Kubik et al., 2007). In a novel environment, animals need to acquire a new spatial representation. Learning of a new spatial map likely requires plastic changes of the hippocampal circuitry and thus high levels of IEG expression. However, at present it is not clear if this activation/plasticity of the hippocampus is induced by the mere visual input from the novel environment or if it requires active exploration of the environment. If hippocampal IEGs expression is related to the activation of spatially responsive cells (place cells, boundary cells etc., Ben-Yishay et al., 2021; Payne at al., 2021), active exploration of a novel environment should cause a higher number of c-Fos expressing cells. Whenever a new location is visited during the exploration, the cells that encode the corresponding location should increase their firing rate, inducing c-Fos expression. The more locations are visited the higher number of hippocampal cells should express c-Fos. Moreover, based on previous studies carried out in other bird species, one can expect to find a higher degree of activation in the anterior segment of Hp during spatial mapping (Payne et al., 2021). However, no study so far reported differences in activation between the anterior, intermediate, and posterior hippocampal segments in domestic chicks.

This study aimed to investigate the impact of active exploration on hippocampus of young domestic chicks, during formation of a new spatial map. To trigger hippocampal activation, we exposed chicks to a novel environment. Using c-Fos imaging, we measured the activation of the anterior, intermediate, and posterior segment of hippocampus in chicks exploring a novel environment. This was compared to a control group that observed the novel environment from a restricted area, without exploring it. We expected to find more c-Fos immunoreactive cells after active exploration of the novel environment. Moreover, we expected that this increased activity would be more visible in the anterior segment of chicks’ hippocampus, as observed in titmice, zebra finches, and owls (Agarwal et al., 2021 preprint; Payne et al., 2021). We also measured the activation of nucleus taeniae of amygdala (TnA), which is a homolog of the mammalian medial amygdala (Abelan et al., 2009). Chick’s TnA responded to a novel environment in our previous study (Morandi-Raikova and Mayer, 2020). We hypothesise the presence of a functional connection between this structure and chicks’ hippocampus and thus a similar activation pattern in both structures. As a control region we measured the activity in the intermediate medial mesopallium (IMM), which is involved in filial imprinting (Horn, 2004) and was not responsive to novel environments in our previous studies (Mayer et al., 2018; Morandi-Raikova and Mayer, 2020;).

## METHODS

### Subjects

We used twenty-four male domestic chicks (*Gallus gallus domesticus*) of the Aviagen ROSS 308 strain. The eggs were obtained already fertilized from a commercial hatchery (CRESCENTI Società Agricola S.r.l. –Allevamento Trepola – cod. Allevamento127BS105/2). Incubation and hatching occurred in complete darkness. After hatching, chicks were individually housed in metal cages (28×32×40cm; W x H x L) with food and water ad libitum, at a constant temperature of 30–32 °C and variable light conditions of 14 h light and 10 h dark. Chicks were food deprived 3 hours before the training on post-hatching day 4. During the training in the experimental room (28 °C), chicks received mealworms (*Tenebrio molitor larvae*) as food rewards, while water was always present ad libitum. At the end of the training sessions, all chicks returned to the animal house and remained there until the next day with food and water ad libitum. On post-hatching day 5, all chicks were tested and subsequently perfused. The experiment was carried out in accordance with the ethical guidelines current to European and Italian laws. All the experimental procedures here described were licensed by the Ministero della Salute, Dipartimento Alimenti, Nutrizione e Sanita’ Pubblica Veterinaria (permit number 560/2018-PR).

### Experimental setup

The experimental apparatus was composed of two compartments. During training, the apparatus consisted of a ‘home compartment’ (28 × 40 × 32 cm, W x H x L) and a ‘habituation compartment’ (40 × 31 × 30 cm). The wall that divided the ‘home compartment’ from the ‘habituation compartment’ could slide vertically and reveal an opening (15 cm x 15 cm) in its centre. This door allowed chicks to enter the ‘habituation compartment’ from the ‘home compartment’. During training, the ‘habituation compartment’ was located inside a ‘novel environment compartment’ (60 × 60 × 60 cm), which was however not visible to the chicks (Figure 1A). The inner surfaces of the setup were white. The walls of the ‘habituation compartment’ were composed of a metal grid. On the outer side of the grid additional walls made of white plastic were present. These acted as visual occluders and prevented chicks to see the ‘novel environment compartment’ during the training. During the test either the visual occluders only (Figure 1B) or the visual occluders and the grid (Figure 1C) could be removed. The setup was equipped with a digital camera (Microsoft LifeCam Studio HD 1920 × 1080 p), placed on the top of the apparatus to record chicks’ behaviour.

**Figure 1:**
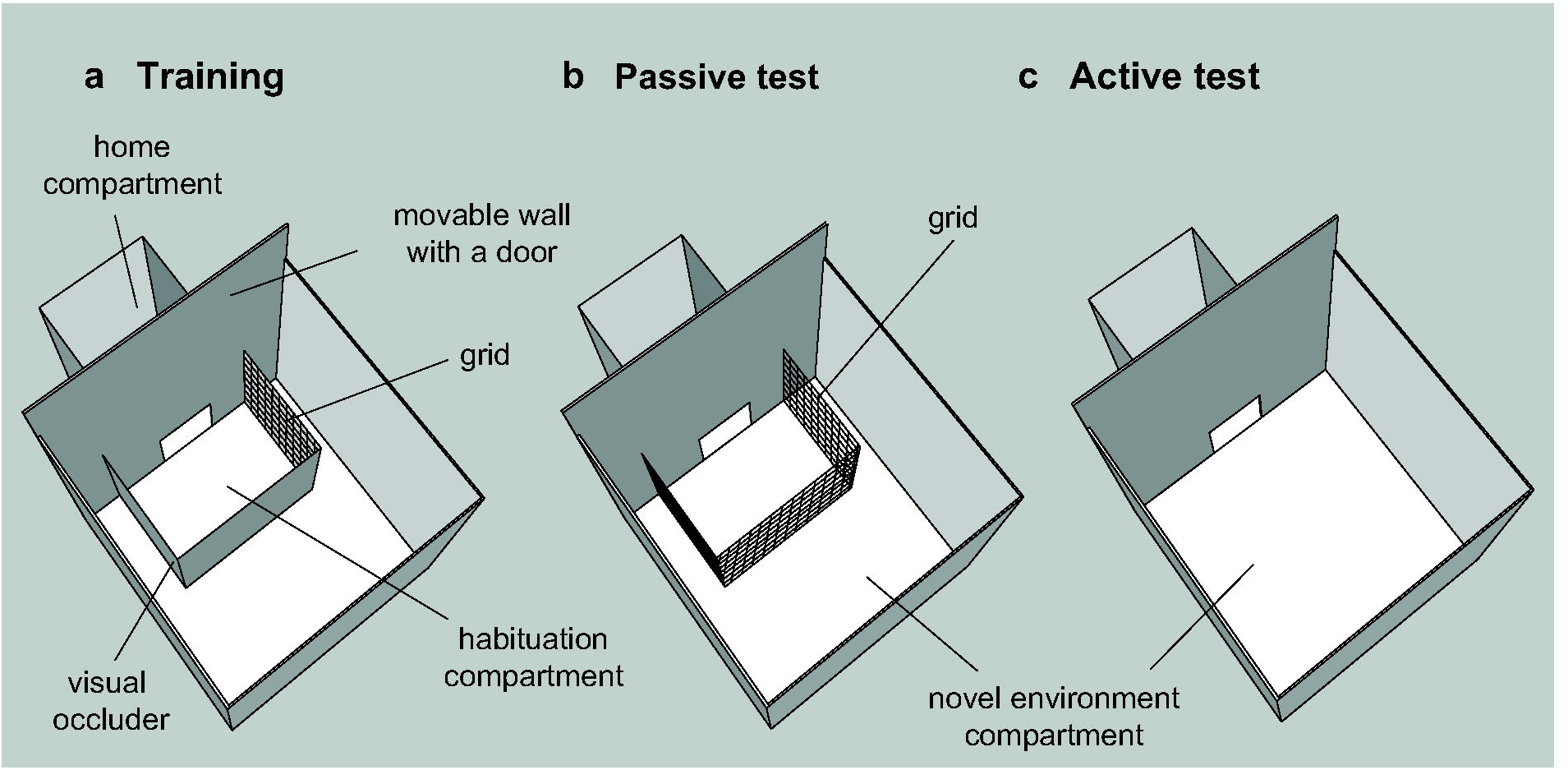
Experimental setup and procedures. **a**) During training the grid with the visual occluder was placed inside the ‘novel environment compartment’. All chicks were trained to walk through the open door and forage for a mealworm in two distinct compartments (‘home compartment’ and ‘habituation compartment’. At test chicks were divided into two different groups. **b**) The ‘passive exploration’ group was tested with the same grid available at training, but this time the visual occluder was removed. Chicks could see the ‘novel environment compartment’, but not explore it actively. **c**) For the ‘active exploration’ group the entire ‘habituation compartment’ was removed and the ‘novel environment compartment’ was exposed. Chicks could enter the ‘novel environment compartment’ and actively explored it.

### Habituation training

All chicks underwent a habituation training on post-hatching day 4. This training aimed to familiarize the animals with the experimental apparatus and to train them to enter and exit the ‘habituation compartment’. For this purpose, domestic chicks were individually placed into the ‘home compartment’ for 30 min of acclimatization, where they received 2 to 3 mealworms. After the acclimatization period, chicks started the training. At the beginning of each training trial, the wall that divided the compartments was lifted, the door appeared and chicks could enter the ‘habituation compartment’ that contained a mealworm. Then, the wall slid down and the door disappeared. After 1 min of permanence in this compartment, the door appeared again and chicks returned to the ‘home compartment’, which contained a mealworm, remaining in it for an additional 1 min. This procedure was repeated ten times for each training session. Each subject underwent in total 6 training sessions, 3 in the morning and 3 in the afternoon. During the intersession intervals (30 min), chicks remained inside the ‘home compartment’.

### Test session for c-Fos labelling

On post-hatching day 5, chicks were divided into two experimental groups: an ‘active exploration’ group and a ‘passive exploration’ group. Before the test, chicks were taken to the experimental room and placed inside the ‘home compartment’, where they remained undisturbed for 5 h. At test, chicks of the ‘active exploration’ group could enter the ‘novel environment compartment’ and explored it for 1 h (Figure 1C). In contrast, chicks of the ‘passive exploration’ group entered the ‘passive compartment’ (Figure 1B), where they remained for 1h. For this group, only the visual occluders were removed from the ‘habituation compartment’. The ‘novel environment compartment’ was visible through the grid, which restricted chicks from entering it. Thus, chicks belonging to the ‘passive exploration group’ could see the ‘novel environment compartment’, but not explore it actively.

### Immunohistochemical procedure

Immediately after the test, all chicks were overdosed with an intramuscular injection of 0.4 ml of 1:1 xylazine (2 mg/ml) and ketamine (10 mg/ml) solution. For visualizing the immediate early gene product c-Fos, brains were processed blind to the experimental conditions and a standard immunohistochemical protocol adapted to chicks was used.

Chicks were perfused transcardially with cold phosphate-buffered solution (PBS; 0.1□mol, pH = 7.4, 0.9% NaCl, 4 °C) and paraformaldehyde (4%PFA in PBS, 4 °C). The heads were severed from the body and placed for 7 days into a 4% PFA/PBS solution for post-fixation. Brains were extracted from the skulls with the use of a stereotaxic head holder (Stoelting). To ensure that the subsequent coronal brain sections would have the same orientation as described for chick’s brain atlas (Kuenzel and Masson, 1988) the horizontal axis of the skulls was oriented at 45° in respect to the horizontal axis of the stereotaxic apparatus.

Brain hemispheres were then separated and embedded into gelatine (7%) containing egg yellow. After an incubation in 20% sucrose in 4% PFA/PBS for 48□h and further 48□h in 30% sucrose in 0.4% PFA/PBS at 4□°C sections were cut. Brains were cut and frozen with the use of a cryostat (Leica CM1850 UV). During cutting four series of 40 μm coronal sections containing the regions of interest (corresponding to one third of the most posterior part of the telencephalon) were collected. Only sections of the first series were used for labelling, while the others were stored as backups.

Endogenous peroxidase activity was depleted with 0.3% peroxide in PBS for 20 min. The sections were then treated with 3% normal goat serum (S-1000; Vector Laboratories, Burlingame, CA, USA) in PBS for 30 min at room temperature to block unspecific binding sites. Anti-c-Fos antibody solution (1:1500 in PBS; rabbit polyclonal, AF-488, Santa Cruz, CA, USA) was applied for 48□h at 4□°C. Afterward, all brain sections were transferred into the secondary antibody solution (1:200 in PBS; biotinylated anti-rabbit made in goat, BA-1000 Vector Laboratories) for 60 min at room temperature. The ABC kit (Vectastain Elite ABC Kit, PK 6100; Vector Laboratories) was used for signal amplification, this step was followed by visualization with VIP kit (SK-4600; Vector Laboratories). Lastly, all sections were mounted on gelatine-coated slides, dried (50□°C), counterstained with methyl green (H-3402; Vector Laboratories) and cover-slipped with Eukitt (FLUKA).

### Brain analysis

Brains were examined blind to the experimental groups and hemispheres with a Zeiss microscope (objective magnification 20×, numerical aperture 0.5; eyepiece 10×), connected to a digital camera (Zeiss AxioCam MRc5) and a computer with the imaging software ZEN. For the analysis, a standard rectangular counting area (150 ×250 µm) was positioned, within the regions of interest, over the spots with the highest number of c-Fos-ir cells. When positioning the counting area, a minimum distance of 20 µm to the borders was always kept. Subsequentially, every activated c-Fos-ir cell was marked with the ZEN software, which then computed the total counts. The measured values derived from different sections were averaged for each area and subsequently standardized to cells/mm^2^.

For the analysis of the hippocampal formation (HF), ten to fifteen sections of each brain hemisphere were used. Based on the anatomical landmarks and its shape, HF was divided into anterior (A 8.6 to A 8.0), intermediate (A 7.8 to A 7.0) and posterior parts (A 6.8 to A 4.6) (Kuenzel and Masson, 1988). In addition, the intermediate and the posterior parts of HF were further subdivided into: ventral (V), dorsomedial (DM) and dorsolateral (DL) (see Figure 2).

**Figure 2:**
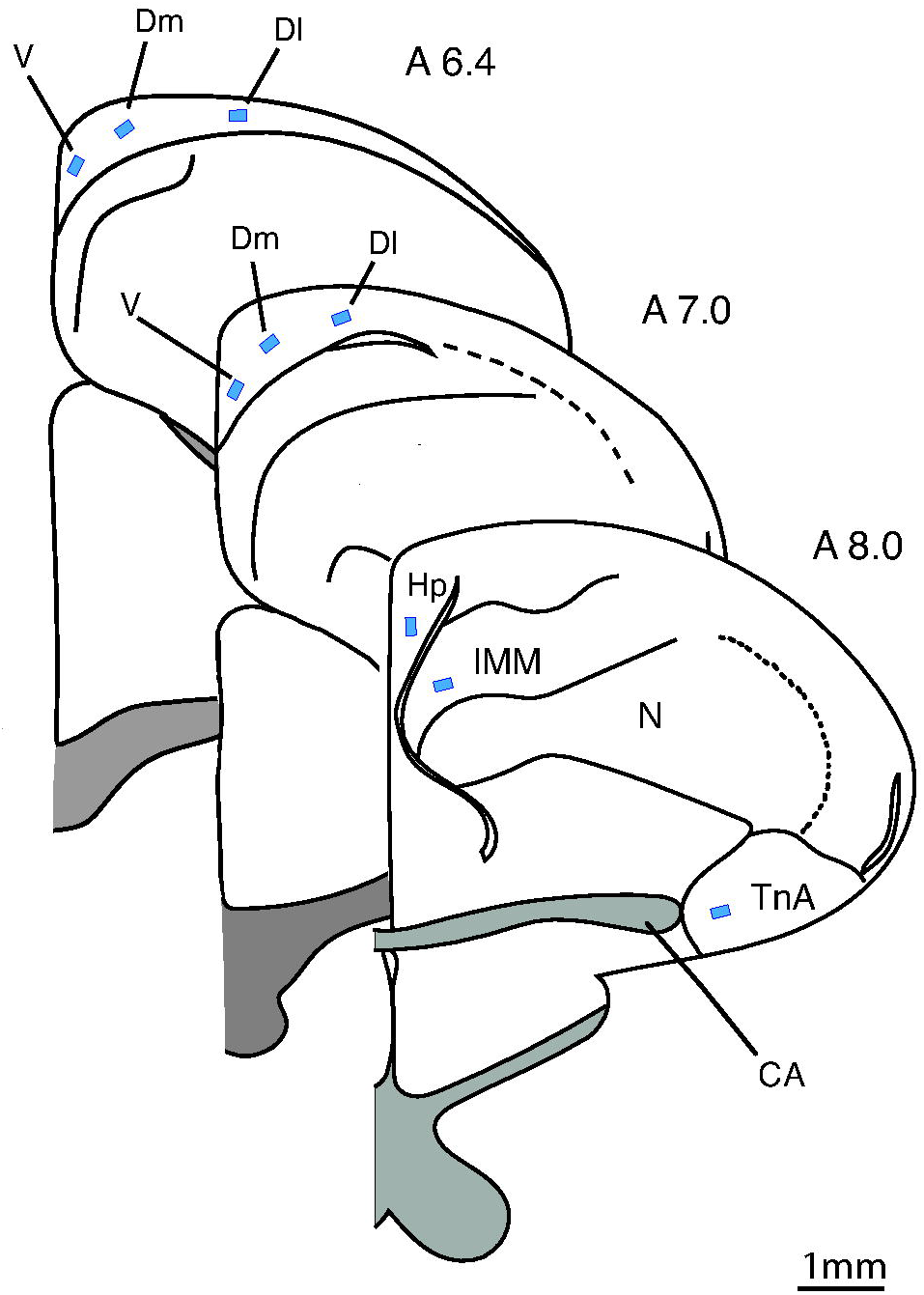
Typical placements of cell counting zones (blue rectangles) in the regions of interest. The intermediate (A 7.0) and posterior (A 6.4) segments of the hippocampal formation were portioned into ventral, dorsomedial and lateral parts. Hp: hippocampus; V: ventral; Dm: dorsomedial; Dl: dorsolateral; IMM: intermediate medial mesopallium; TnA: nucleus taeniae of the amygdala; N: nidopallium; CA: anterior commissure.

To quantify c-Fos-ir cells in nucleus taeniae of the amygdala (TnA), five sections corresponding to the A 7.4 and A 6 .4 of the brain atlas (Kuenzel and Masson, 1988) of both hemispheres were selected. Finally, for the quantification of c-Fos-ir cells contained within IMM, this brain region was outlined according to the drawings of Ambalavanar et al., 1993. The counting in this area was performed on five brain sections from A 8.6 to A 4.6.

### Behavioural analysis

Video recordings of the test session were analysed offline with EthoVision 3.1 (Noldus Information Technology, Leesburg, VA, Noldus et al., 2002). Videos were analysed at a rate of 6-samples/s, while to track animals’ position (x, y coordinates), the background subtraction method was used. These coordinates in px were converted in cm by calibrating the software to the width of the experimental compartments. Behavioural analyses were carried out for both experimental groups. The behavioural parameters that were extracted through EthoVision were: distance moved (cm), velocity (cm/s), number of visited sectors and sector change frequency. To extract these last two behavioural variables, both experimental compartments were subdivided into equal zones of 10 cm x 10 cm, to represent the various locations domestic chicks could have visited at test. Thus, the ‘novel environment compartment’ was subdivided into 36 different sectors (6 × 6, rows x columns) while the ‘passive compartment’ into 12 sectors (3 × 4). By computing the number of sectors visited by each chick, we could measure how widely each animal explored the available space. For instance, a chick of the active exploration group that always remained motionless during the test, would have visited only one sector, while a chick that explored all the available space would have a score of 36 visited sectors. In contrast, the frequency of sector changes and the distance moved by each animal revealed how intensively the animal moved across space, regardless of the absolute number of locations visited. Additionally, to assess whether the motivation to explore the available space differed among groups, a percentage of explored sectors was computed by dividing the number of visited sectors by the overall available sectors for each individual. Data from Ethovision was then exported into the software IBM SPSS Statistics (v. 20) and analysed.

### Statistical analysis

To analyse whether there was any difference in the measured behavioural parameters between the two experimental groups, a multivariate analysis of variance (MANOVA) was conducted. To reveal differences in the activation pattern of the investigated brain areas between the two experimental groups, a repeated-measures ANOVA was performed. This analysis included a between-subject factor ‘groups’ (2 levels: ‘active exploration’ group, ‘passive exploration’ group) and two within-subject factors: ‘area’ (9 levels: anterior Hp, intermediate ventral Hp, intermediate dorsomedial Hp, intermediate dorsolateral Hp, posterior ventral Hp, posterior dorsomedial Hp, posterior dorsolateral Hp, TnA, IMM) and ‘hemispheres’ (2 levels: left, right). Subsequentially, for the post-hoc analyses, an independent samples t-tests was conducted. To assess whether there was any relationship between the measured behavioural parameters and the activation of the investigated brain regions, a Pearson correlation analysis was run.

The alpha level of 0.05 was considered significant, however the obtained values were also tested against Bonferroni-adjusted alpha level of 0.006 (0.05/9). We report F- and t-statistics, exact p-values, means, standard error of the means (s.e.m.) and standardized effect sizes (Cohen’s d for t-tests and □_p_^2^ for ANOVAs). All statistical analyses were performed using SPSS and R (R Core Team, 2020), while the graphs were created with the software GraphPad Prism 8 and R. Datasets used for the statistical analysis are in supplement (Table S1).

## RESULTS

### Behavioural results

Analyses of chick’s walking tracks revealed significant differences between the groups (Figure 3). As expected, chicks of the ‘active’ exploration group moved longer distances compared to the ‘passive’ group (‘active’: 4013.72±565 cm; ‘passive’: 1754.48±259 cm; F_(1,22)=_13.232, p=0.001, □_p_^2^=0.376, mean±s.e.m. rounded numbers,) and visited more sectors (‘active’: 31.7±2.1; ‘passive’: 9.5±0.7; F_(1,22)=_99.835, p<0.001, □_p_^2^=0.819). They also moved faster (‘active’: 4.44±0.6 cm/s; ‘passive’: 2.61±0.6 cm/s; F_(1,22)=_5.399, p=0.030, □_p_^2^=0.197) and changed more often the sectors during exploration of the novel environment (‘active’: 470.3±7; ‘passive’: 200.1±4; F_(1,22)=_12.450, p=0.003, □ ^2^=0.361). However, the percentage of available space covered by chicks at test was not different between the two groups (‘active’: 87.96±5.8 %; ‘passive’: 79.17±6.1 %; F_(1,22)=_1.102, p=0.305, □_p_^2^=0.048).

**Figure 3:**
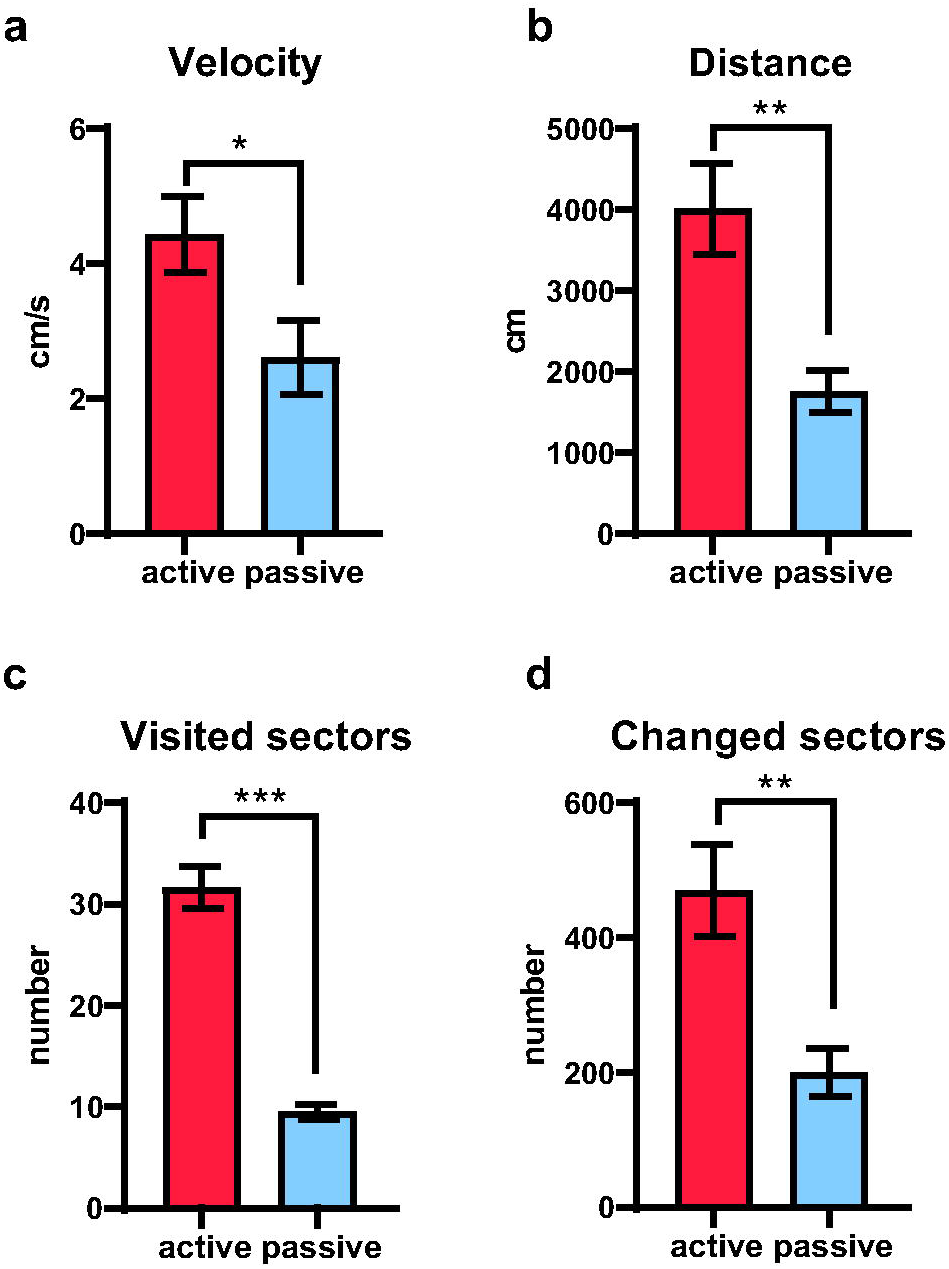
Behavioural performance during test of the ‘active exploration’ group in red and ‘passive exploration’ group in blue. Chicks of the ‘active’ exploration group moved faster (a), longer distances (b), visited more sectors (c) and changed more often the sectors (d) compared to the ‘passive’ group. Bar plots show mean and s.e.m. (*p<0.05, **p<0.01).

### Brain results

All twenty-four brains (n=12 per each experimental group) were successfully stained for c-Fos. The nuclei of c-Fos immunoreactive (-ir) cells were stained black and were easily discernible from the other cells counterstained with methyl green (Figure 4). Measured c-Fos-ir cells densities are summarized in Table 1.

**Figure 4:**
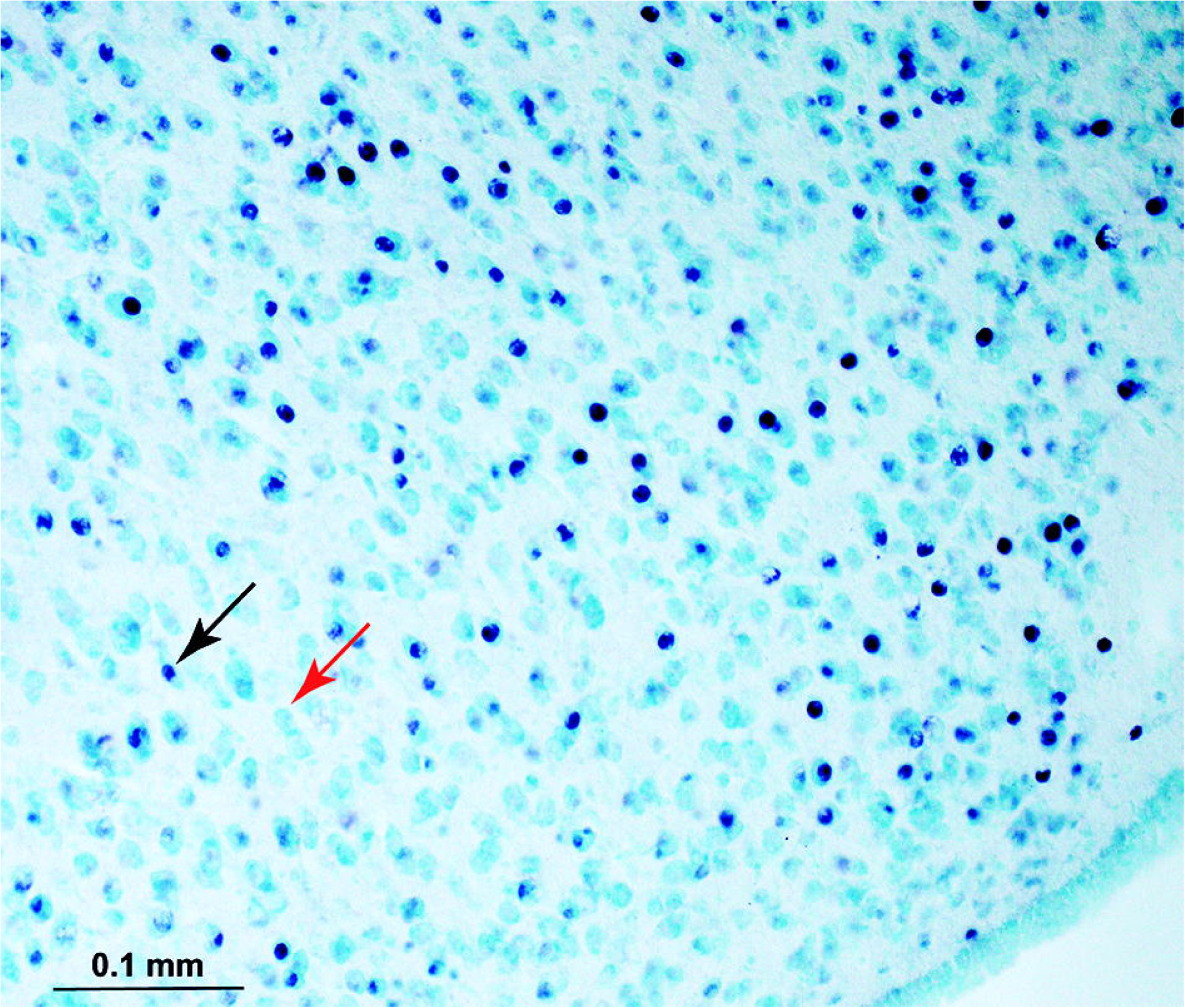
Example photo of a hippocampal section of an experimental chicks. c-Fos-ir (immunoreactive) cells are stained black after a successful immunohistochemical procedure (black arrow) and are easily discernible from the methyl-green counterstained cells (red arrow).

**Table 1:** Measured cell densities (c-Fos-ir cells/mm^2^) across brain areas of the left and right hemispheres and total values after the two hemispheres were lumped together (mean□±□s.e.m., rounded numbers). Hp: hippocampus; int: intermediate; V: ventral; DM: dorsomedial; DL: dorsolateral: post: posterior: TnA: nucleus taeniae of the amygdala; IMM: intermediate medial mesopallium.

|  | Active exploration group |  |  | Passive exploration group |  |  |
| --- | --- | --- | --- | --- | --- | --- |
|  | Left | Right | Total | Left | Right | Total |
| **Anterior Hp** | 1788.9±185 | 1670.4±197 | 1729.6±164 | 998.5±148 | 1094±188 | 1046.3±142 |
| **Int. V Hp** | 320.2±57 | 408.6±65 | 364.4±46 | 296.8±65 | 274.1±45 | 285.4±44 |
| **Int. DM Hp** | 688.3±127 | 802.3±126 | 745.3±106 | 606.6±104 | 538.1±97 | 572.4±90 |
| **Int. DL Hp** | 1937±181 | 2054.4±177 | 1995.7±139 | 1372.3±221 | 1554.4±211 | 1463.4±162 |
| **Pos. V Hp** | 233.3±41 | 313.3±44 | 273.3±38 | 273±53 | 252.6±59 | 262.8±50 |
| **Pos. DM Hp** | 492.6±64 | 544.8±72 | 518.7±61 | 484.1±67 | 518.2±123 | 501.1±79 |
| **Pos. DL Hp** | 1927±294 | 2304.8±212 | 2115.9±213 | 1828.9±235 | 1828.9±251 | 1828.9±168 |
| **TnA** | 1534.4±155 | 1561.1±201 | 1547.8±160 | 921.1±100 | 953.3±82 | 937.2±68 |
| **IMM** | 1165.8±157 | 1020±142 | 1092.9±137 | 1024.5±149 | 1048±117 | 1036.2±105 |

The repeated measures ANOVA revealed a significant interaction of ‘area’ and ‘group’ (p=0.001). This indicates that c-Fos-ir cell densities were different between the two experimental groups, in an area-specific fashion. However, there was no effect of ‘hemisphere’ nor any interaction between ‘hemisphere’ and ‘area’ or ‘group’ (see Table 2 for all ANOVA results). Therefore, for the post hoc analysis the measures for the left and right hemispheres were lumped together.

**Table 2:**
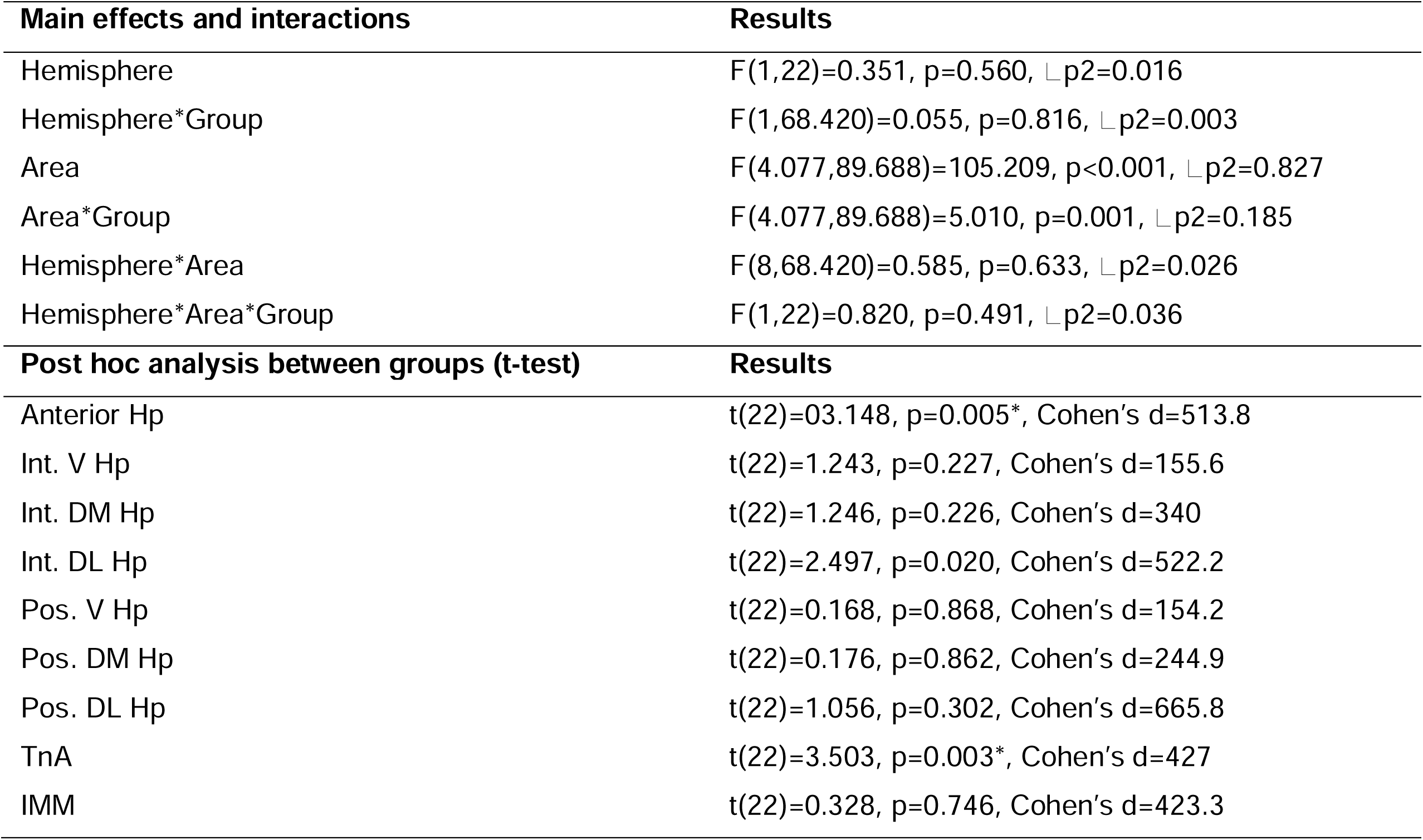
Results of the repeated measures ANOVA and the corresponding post hoc analysis. **\*** Significant also after a Bonferroni adjustment for multiple comparisons.

| Main effects and interactions | Results |
| --- | --- |
| Hemisphere | $F(1,22)=0.351$, $p=0.560$, $\eta^2_p=0.016$ |
| Hemisphere*Group | $F(1,68.420)=0.055$, $p=0.816$, $\eta^2_p=0.003$ |
| Area | $F(4.077,89.688)=105.209$, $p<0.001$, $\eta^2_p=0.827$ |
| Area*Group | $F(4.077,89.688)=5.010$, $p=0.001$, $\eta^2_p=0.185$ |
| Hemisphere*Area | $F(8,68.420)=0.585$, $p=0.633$, $\eta^2_p=0.026$ |
| Hemisphere*Area*Group | $F(1,22)=0.820$, $p=0.491$, $\eta^2_p=0.036$ |
| Post hoc analysis between groups (t-test) | Results |
| Anterior Hp | $t(22)=03.148$, $p=0.005^*$, Cohen's $d=513.8$ |
| Int. V Hp | $t(22)=1.243$, $p=0.227$, Cohen's $d=155.6$ |
| Int. DM Hp | $t(22)=1.246$, $p=0.226$, Cohen's $d=340$ |
| Int. DL Hp | $t(22)=2.497$, $p=0.020$, Cohen's $d=522.2$ |
| Pos. V Hp | $t(22)=0.168$, $p=0.868$, Cohen's $d=154.2$ |
| Pos. DM Hp | $t(22)=0.176$, $p=0.862$, Cohen's $d=244.9$ |
| Pos. DL Hp | $t(22)=1.056$, $p=0.302$, Cohen's $d=665.8$ |
| TnA | $t(22)=3.503$, $p=0.003^*$, Cohen's $d=427$ |
| IMM | $t(22)=0.328$, $p=0.746$, Cohen's $d=423.3$ |

The post hoc analysis revealed significantly higher c-Fos-ir cell densities for the ‘active exploration’ group compared to the ‘passive exploration’ group in the anterior Hp (p=0.005, significant also after the Bonferroni correction: p=0.045), in the intermediate DL Hp (p=0.020) and in TnA (p=0.003, significant also after the Bonferroni correction: p=0.027). No significant difference between the two experimental groups were found in the other hippocampal subdivisions and in the IMM (Figure 5, Table 2).

**Figure 5:**
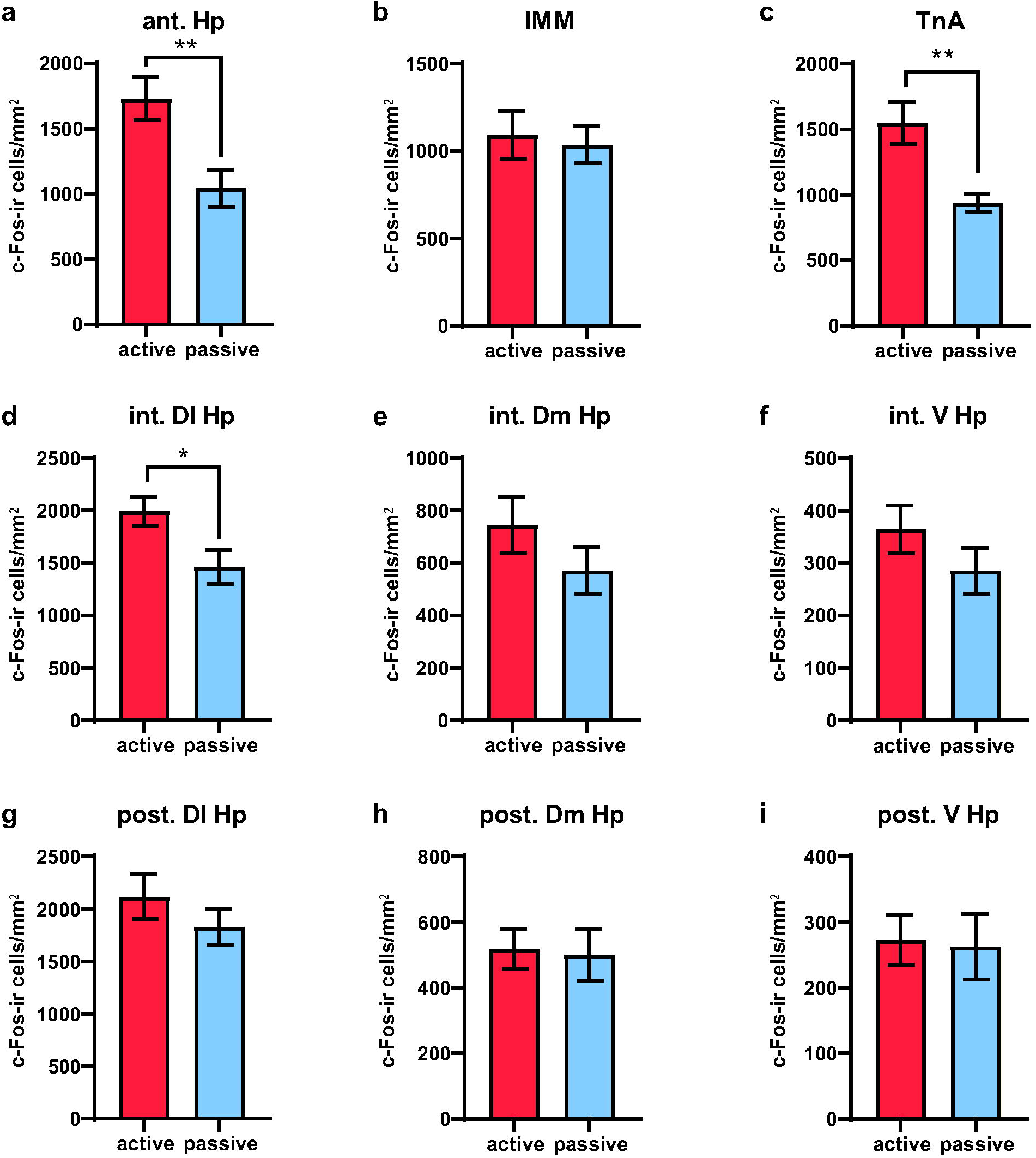
Measured c-Fos-ir (immunoreactive cell densities) across brain regions of the two experimental groups. **a**) Anterior hippocampus (ant. Hp). **b**) intermediate medial mesopallium (IMM). **c**) Nucleus taeniae of the amygdala (TnA). d) intermediate ventral hippocampus (int. V Hp). **e**) Intermediate dorsomedial hippocampus (int. Dm Hp). **f**) Intermediate dorsolateral hippocampus (int. Dl Hp). **g**) Posterior ventral hippocampus (post. V Hp). **h**) Posterior dorsomedial hippocampus (post. Dm Hp). **i**) Posterior dorsolateral hippocampus (post. Dl Hp). Bar plots show mean and s.e.m. (*p<0.05, **p<0.01).

### Correlations

A significant correlation was found between the number of visited sectors and the density of c-Fos-ir cells in the anterior Hp (r=0.489, p=0.016) (Figure 6B). Furthermore, also the density of c-Fos-ir cells in the TnA correlated with the number of visited sectors (r=0.447, p=0.030) (Figure 6C). No correlation between number of visited sectors and c-Fos-ir cell density was found in the other hippocampal subdivisions nor in IMM. Moreover, no correlation was found between the other behavioral parameters (velocity, distance moved and sector change frequency) and the density of c-Fos expression in any of the investigated brain areas (see Figure 6A for all Pearson correlation results).

**Figure 6:**
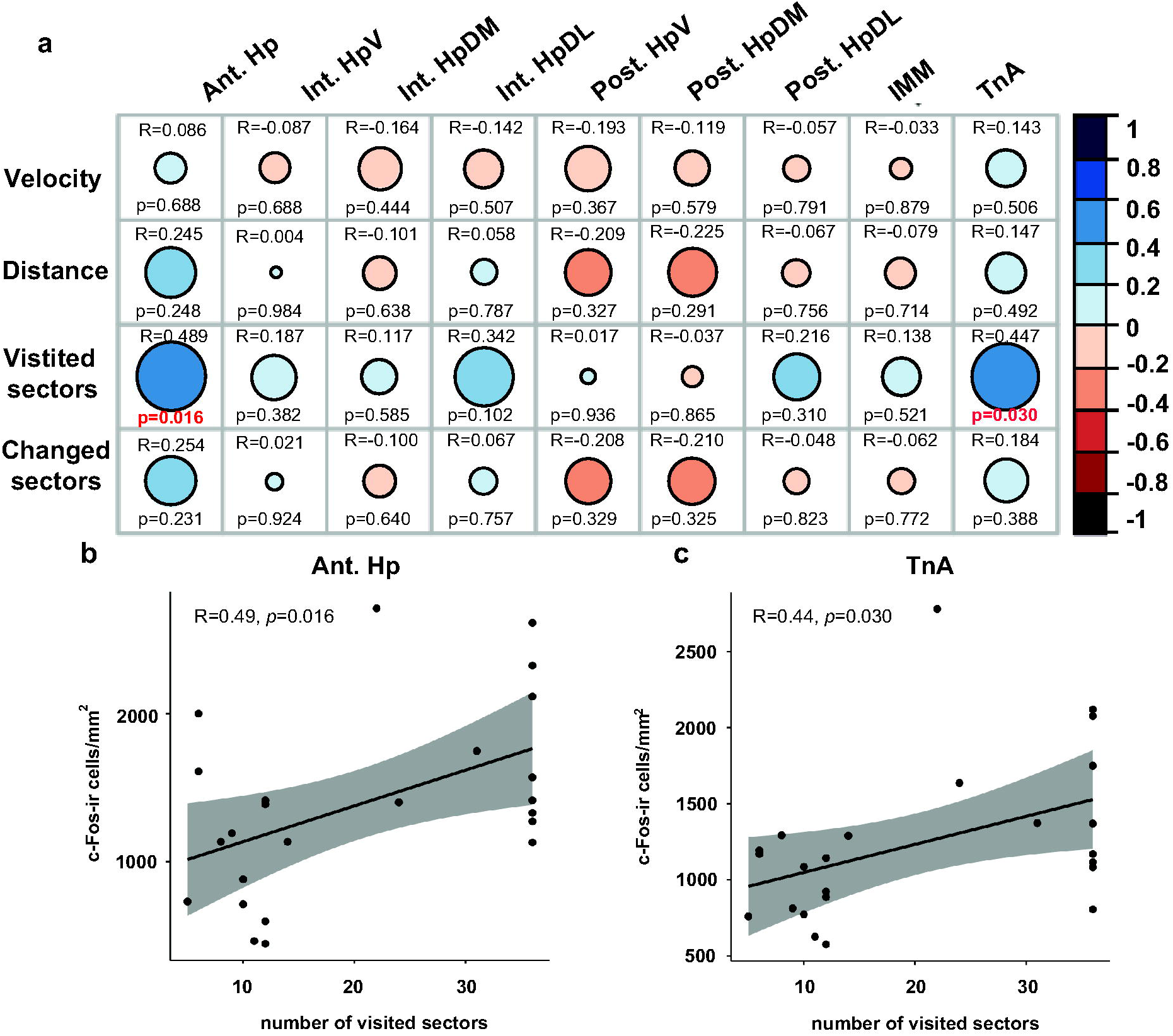
Brain-behaviour correlations. (**a**) Correlation matrix of all brain regions and behavioural parameters. The size of the dots corresponds to the strength of correlation while the colour to the direction (blue=positive, red=negative). (b and c) Significant correlations between visited sectors and density of c-Fos-ir cells within the anterior hippocampus (Ant. Hp) and nucleus taenia of the amygdala (TnA).

## DISCUSSION

Domestic chicks that could actively explore a novel environment had a higher c-Fos expression in the anterior hippocampus and in the dorsolateral parts of the intermediate hippocampus. Furthermore, a higher c-Fos expression was observed in TnA of the ‘active exploration’ group, compared to the group confined to a restricted area. The difference was region-specific. In IMM and in the posterior hippocampal segment, c-Fos activity did not differ between the ‘active’ and ‘passive’ exploration groups.

Our results confirm that the avian hippocampus responds to a novel environment, as observed in mammals (Kubik et al., 2007). This is in line with our previous studies showing hippocampal sensitivity to environmental novelties, such as size and shape changes, in domestic chicks (Mayer et al., 2018; Morandi-Raikova and Mayer, 2020). However, here we show for the first time that hippocampus activation is higher only in chicks that actively explored the novel environment. Thus, a purely visual input is not sufficient to trigger the full extent of the hippocampal activation in response to the novel environment. As we predicted, physical movement across different locations crucially contributed to hippocampal activation. Indeed, the number of visited sectors positively correlated with the activation in the anterior hippocampus. Overall, chicks of the ‘active exploration’ group moved more, faster, they changed more frequently the sectors and explored more sectors than the chicks of the ‘passive exploration’ group. These results are not surprising, since chicks of the ‘active exploration’ group could move within a larger area, compared to the chicks of the ‘passive exploration’ group. However, both groups covered overall a similar percentage of available space, meaning that their motivation to explore did not differ. Furthermore, the behavioural parameters ‘distance moved’, ‘velocity’ and the frequency of ‘sector changes’ did not correlate with brain activity in any of the regions of interest. Thus, the brain activity in the anterior, dorsolateral intermediate Hp, and TnA was not influenced by, how much or how fast the animals moved, but only by how many spatial locations they visited overall.

These findings suggest that, in birds like in mammals, the increase in hippocampal c-Fos expression during exploration of a novel environment reflects the increased firing rates of spatially coding neurons. In mammals, different hippocampal cells encode different locations and other properties of the allocentric environment (Poulter et al., 2018). The more place fields the animal crosses, the larger the number of different place cells that are activated (O’Keefe and Nadel, 1978; Poulter et al., 2018). Given that neural activation induces immediate early gene expression, we predicted that the number of explored fields would positively correlate with the density of c-Fos expressing cells, which we indeed found in our study. Furthermore, immediate early gene expression is typically linked to learning related plasticity (Jones et al., 2001; Guzowski, 2002; Barry and Commins, 2011). In a novel environment, a new spatial map needs to be formed. For instance, spatially coding cells need to remap to new firing fields (Muller and Kubie, 1987; Fyhn et al., 2007; Barry et al., 2012) and new environmental boundaries need to be represented (Sargolini et al., 2006).

The newly formed spatial map needs to be memorized, leading to plastic changes of the hippocampal network (Sheng and Greenberg, 1990; Lanahan and Worley, 1998; Kubik et al., 2007). The structural changes to the neurons are mediated through the expression of immediate early gene products, including c-Fos (Sheng and Greenberg, 1990; Lanahan and Worley, 1998). Therefore, exploration of a novel environment should induce high levels of hippocampal IEG expression, as we observed here (Kubik et al., 2007).

The increased c-Fos expression, in the anterior and the intermediate dorsolateral hippocampal parts of the ‘active exploration’ group, further supports the idea that spatially coding cells exist in domestic chicks’ hippocampi. Our findings are particularly interesting in the light of the divergent results coming from studies on single-unit activity in the hippocampus of different bird species. The recent discovery of place cells in the anterior hippocampi of two Neoaves, titmice and zebra finches (Payne et al., 2021), may indicate that this function was probably already present in the common ancestor of sauropsids and mammals. However, a recent study of hippocampal activity in a galliform specie, quail (*Coturnix coturnix*), failed to identify place cells. In this study, many hippocampal neurons were successfully recorded, but only head-direction cells were identified among them (Ben-Yishay et al., 2021). This evidence may be used to argue that place cells would exist only in some bird species while in others not. For instance, one could hypothesise that place cells represent a case of convergent evolution between mammals and the most cognitively advanced birds, such as Neoaves, absent in Galliformes. In this view, Indeed, it has been argued that Galliformes, such as quails and chickens, retained more ancestral traits compared to Neoaves (Prum et al., 2015). In contrast, our results with c-Fos indirectly suggest that place cells (and other spatially coding cells) may exist also in Galliformes. Likely that the density of spatially coding cells is much lower in Galliformes compared to Neoaves, making them more difficult to detect electrophysiologically. Indeed, the number of spatially coding neurons vary in different bird species, probably due to divergent ecological adaptations. For example, fewer place neurons could be found in the hippocampus of a non-food hoarding species (zebra finches), compared to the food-hoarding titmice (Payne et al., 2021). Even fewer of such neurons may exist in domestic chicks and quails, in line with the presence of more ancestral traits in these birds (Prum et al., 2015).

It is important to also consider that our current results do not provide information on the specific neural populations that were activated. In addition to place cells and head directions cells, other spatially coding cells may contributed to the effect in the ‘active’ group (e.g., border cells or grid cells, if existent in birds’ hippocampi). Electrophysiological confirmation studies with domestic chicks are urgently needed at this point. Our results indicate that, in such studies, recordings should be performed in the anterior hippocampus (in line with Payne et al., 2021). It is also worth mentioning that subdivisional differences in c-Fos expression in chick’s hippocampus have been shown in our earlier studies too. For instance, social novelties activated the ventral and dorsomedial parts, but not the dorsolateral parts, of the intermediate hippocampus (Corrales-Parada et al., 2021). On the contrary, here we found significant activation of the dorsolateral intermediate hippocampus and the anterior hippocampus. This highlights a potential segregation of social and spatial functions between areas of the ventral and dorsal avian hippocampus, which needs further investigation.

Functional differences between the hippocampi of the two hemispheres have been reported in earlier studies with chicks (Tommasi et al., 2003; Corrales-Parada et al., 2021, Morandi-Raikova and Mayer, 2021). For instance, when we trained chicks to navigate in a large arena in relation to free-standing objects, only the right hippocampus showed c-Fos upregulation (Morandi-Raikova and Mayer, 2021). In contrast, no lateralization was found in the present study. This discrepancy may be explained by the different nature of the two tasks. In our previous study chicks were specifically trained to orient using the relational information provided by free-standing objects (Morandi-Raikova and Mayer, 2021). The results suggested that spatial-relational computations are predominantly processed by the hippocampus of the right hemisphere. On the contrary, in the present study, chicks had to explore a novel environment and to acquire a new spatial map based on the geometrical shape of the environmental layout. This task is less specific, since multiple strategies can be used to orient in this situation. Thus, the task may have activated multiple hippocampal functions, which are processed in parallel in both hemispheres during the acquisition of a new spatial map. This lack of lateralization is in line with our previous studies, where chicks had to orient by the geometrical layout of the environment (Mayer et al., 2016), or were simply exposed to novel environmental shapes (Mayer et al., 2018; Morandi-Raikova and Mayer, 2020). In none of these studies we were able to detect clear lateralization. More research using different orientation tasks is needed to characterize the functional specializations of the left and the right hippocampi in birds.

With the present study we confirm that, in birds like in mammals, exposure to novel environments induces hippocampal activation, which is likely related to the formation of new spatial representations. To best of our knowledge, so far only two other studies investigated the involvement of the avian hippocampus in the exploration of novel environments (Mayer et al., 2018; Morandi-Raikova and Mayer, 2020). Altogether, these studies contribute to the vast literature showing that, despite the fundamental differences in the structure of the hippocampal formation in birds and mammals (Striedter, 2016), its involvement in spatial functions is shared among the two clades (Bingman et al., 2005; Siegel et al., 2005; Hough and Bingman, 2008; Kahn and Bingman, 2009; Mayer et al., 2010, 2013, 2016; Mayer and Bischof, 2013; Coppola et al., 2015, 2016; Sherry et al., 2017; Lormat et al., 2020; Morandi-Raikova and Mayer, 2021; Payne et al., 2021).

Finally, in the present study, we also found higher expression of c-Fos in the nucleus taeniae of the amygdala (TnA) of the ‘active exploration’ group, compared to the ‘passive exploration’ group. This is in line with our previous finding that TnA and the hippocampus activate in chicks exposed to a novel environment (Morandi-Raikova and Mayer, 2021). In addition, in the present study the activation of both taeniae and hippocampus positively correlated with the number of explored sectors. This suggests a functional linkage between these two brain regions, which may play a role for spatial memory formation. This is not unlikely, given that the nucleus taeniae of the amygdala is anatomically interconnected with the hippocampal formation (Casini et al., 1986; Cheng et al., 1999). Furthermore, a functional linkage between the amygdala and the hippocampus has been reported in mammals too (Sheth et al., 2008). Disruption of amygdala activity prevents the increase of hippocampal Fos expression in response to novel environments (Sheth et al., 2008). Based on these findings, it has been proposed that the amygdala may affect hippocampal encoding of specific environmental features. Our findings suggest that in birds, like in mammals, the amygdala may modulate spatial information processing in the hippocampus, which may be a conserved function of the hippocampus-amygdala complex in vertebrates. Whether the role of the amygdala is to encode specific environmental features remains indecisive with the current data. Also, a presence of place cells in the amygdala would be a rather surprising trait, which, as far as we know, has never been investigated in any vertebrate species. An alternative possibility would be to consider that the activity of taeniae might reflect neophobia induced by the novelty itself, as suggested by other studies (Morandi-Raikova and Mayer, 2021; Perez et al., 2021). The more sectors are explored, the more the animal was in the novel open field, which may have induced a stronger neophobic reaction. Alternatively one could speculate on the role of taeniae in processing reward information during acquisition of spatial memories. A reward function of mammalian medial amygdala (homolog of birds taeniae) has been reported in mice at least in a social context (Hu et al., 2021). In summary, the role of taeniae in processing spatial novelties remains speculative at this stage. However, the present study and our previous work (Morandi-Raikova and Mayer, 2021) strongly support the idea that the hippocampus-amygdala complex exists in birds too. Moreover, like in mammals, also in avian species, both structures might play a role in spatial functions.

To conclude, our findings suggest the existence of spatially coding neurons (such as place cells) in the domestic chicken’s anterior hippocampus. IEG products do not represent a direct approach to investigate place cells. However, they provide a great opportunity to image the activation of large populations of cells, which can be aligned with anatomical subdivisions of the hippocampus. Future electrophysiological studies should target the anterior and the dorsolateral portion of the intermediate hippocampus, which increased the density of c-Fos immunoreactive cells after active exploration of a novel environment. Finally, our study suggests a functional linkage of the hippocampus and nucleus taeniae of the amygdala for the processing of spatial information. Many unanswered questions still remain, but the present study opens new doors to study the evolution of the neural circuits behind animal navigation.

## LIST OF ABBREVIATIONS

Hp: Hippocampus
HF: Hippocampal formation
V: ventral
DM: dorsomedial
DL: dorsolateral
Int: intermediate
Post: posterior
TnA: Nucleus taeniae of the amygdala
IMM: Intermediate medial mesopallium
IEGs: Immediate early genes
c-Fos-ir: c-Fos immunoreactive
PBS: phosphate buffered saline
PFA: paraformaldehyde

## ACKNOWLEDGEMENTS

Our special thanks is due to Marta Rodriguez Armenia, who helped with collecting part of the behavioral data and brain processing. Special thanks also to Orsola Rosa-Salva for her useful comments on the manuscript.

## COMPETING INTERESSTS

The authors declare that they have no known competing financial interests or personal relationships that could have appeared to influence the work reported in this paper.

## AUTHOR CONTRIBUTIONS

AMR and UM: conceived and designed the experiments; AMR performed the experiments, analysed the data and together with UM wrote the manuscript; UM supervised the project.

